# Proteomic signature and mesenchymal stromal/stem cell-derived extracellular vesicle treatment of iPSC motor neurons from different ALS patient groups

**DOI:** 10.1101/2022.07.05.498816

**Authors:** Suzy Varderidou-Minasian, Channa E. Jakobs, Svetlana Pasteuning-Vuhman, Annabel Timmers, Maarten Altelaar, Magdalena J. Lorenowicz, R. Jeroen Pasterkamp

## Abstract

Motor neurons (MNs) derived from human induced pluripotent stem cells (iPSCs) offer a powerful model to study motor neuron diseases, such as amyotrophic lateral sclerosis (ALS). While widely used, our knowledge of the proteomic changes in these models is rather rudimentary. In this study, we conducted a comparative proteomic analysis of iPSC-derived MNs carrying ALS-associated mutations in *C9ORF72*, *TARDBP*, or *FUS*. This revealed both mutation-specific and shared proteomic signatures, unveiling common and divergent disease mechanisms. Using these new insights, we then evaluated the therapeutic potential of extracellular vesicles from mesenchymal stromal/stem cells (MSC-EVs). These experiments showed a functional effect of MSC-EVs in ALS-FUS MNs *in vitro* and their ability to reverse proteomic changes more generally in MNs with different ALS genetic backgrounds. These findings highlight key molecular pathways involved in ALS at the protein level and support the potential of MSC-EVs as a versatile therapeutic approach.

**Highlights:**

- This study determines ALS-associated proteomic signatures in iPSC motor neurons (MNs)
- Proteomic signatures indicate common and divergent disease mechanisms in MNs
- MSC-EV treatment restores neurite growth defects in FUS-ALS MNs
- MSC-EVs restore proteomic changes in MNs carrying different ALS mutations

## Introduction

Amyotrophic lateral sclerosis (ALS) is an adult-onset neurodegenerative disorder characterized by progressive muscle weakness and degeneration of motor neurons (MN) in brain and spinal cord (Brown & Al-Chalabi, 2017; van Es et al., 2017). ALS patients eventually die due to respiratory failure within 2 to 4 years following symptom onset. The disease arises from a complex interplay between environmental and genetic factors. Over 40 distinct genes have been linked to ALS (Al-Chalabi et al., 2017) including *Chromosome 9 open reading frame 72* (C9ORF72; ∼40%) (DeJesus-Hernandez et al., 2011; Renton et al., 2011), *Fused in sarcoma* (FUS; ∼1-5%) (Kwiatkowski et al., 2009), and *TAR DNA binding protein 43* (TDP-43; ∼1-5%) (Neumann et al., 2006). Analysis of these genes has provided invaluable insight into the mechanisms underlying ALS, but translation of these findings into effective therapies has been challenging. Drugs currently approved for the treatment of ALS are either modestly effective (e.g. Riluzole) (Traynor et al., 2003; Wei et al., 2024) or only target a small subset of patients (e.g. Tofersen) (Miller et al., 2020). Nevertheless, their development underscores our ability to design therapeutic approaches based on knowledge of disease pathways and indicates that further insight into ALS pathogenesis is needed. Human induced pluripotent stem cell (iPSC)-derived MNs cultures are a powerful tool for dissecting ALS pathogenesis (Pasteuning-Vuhman et al., 2021). Different ALS mutations have been modelled in iPSC-derived MNs (Burkhardt et al., 2013; Chen et al., 2014; N. Egawa et al., 2012; Guo et al., 2017b; Hawrot et al., 2020; Wainger & Lagier-Tourenne, 2018), but their characterization at the proteomic level is rather incomplete. Further, models for different mutations have mostly been studied in isolation rather than being compared to identify both mutation-specific and converging disease pathways. Therefore, in this study we, for the first time, determined and compared proteomic signatures associated with three major ALS mutations (in *C9ORF72*, *FUS* and *TARDBP*) in iPSC-derived MNs. These newly acquired data were subsequently exploited to understand the proteomic mechanisms underlying the effects of treatment with extracellular vesicles (EVs) derived from mesenchymal stromal/stem cells (MSCs) in ALS iPSC-derived MN cultures.

Several studies have demonstrated beneficial effects of MSCs in murine models of ALS (Boucherie et al., 2009; de Munter et al., 2020; Marconi et al., 2013; Martínez-Muriana et al., 2020; Spejo et al., 2018). MSCs have been shown to delay MN degeneration, reduce inflammatory responses, and prolong survival. Emerging evidence suggests that these therapeutic benefits may primarily derive from EVs released by MSCs. EVs represent a heterogeneous group of membrane-bound vesicles that transport diverse cargo, including proteins, RNA, lipids and carbohydrates, to regulate cell development, differentiation and survival (Collino et al., 2010; Varderidou-Minasian & Lorenowicz, 2020). For example, MSC-EVs selectively promote neurite outgrowth in cortical neuron cultures and modulate neural plasticity (Lopez-Verrilli et al., 2016; Xin et al., 2012). To further define the mechanism-of-action of MSC-EVs in the context of ALS, we treated mutant ALS iPSC-derived MNs with MSC-EVs and studied cellular and proteomic changes.

In this study, we determine and integrate the altered proteomic profiles of MNs carrying different ALS-associated mutations. This not only provides unique insights into the molecular disease mechanisms triggered by specific mutations but also highlights key shared molecular pathways that underlie ALS more generally and may act as a starting point for therapy development. For example, our study shows the beneficial effect of MSC-EV treatment in ALS-FUS MNs *in vitro* and their ability to reverse proteomic changes more generally in MNs with different ALS genetic backgrounds. These data support the exciting idea that MSC-EVs may act as therapeutic agents in ALS.

## Results

### Proteomics analysis of iPSC-derived MNs with different ALS backgrounds

To study the effect of ALS-associated gene mutations at the protein level, human MNs were generated from previously reported iPSCs carrying genetic defects in *C9ORF72,* the most common genetic cause of ALS, or carrying ALS-associated mutations in the RNA-binding proteins FUS or TDP-43. Neuronal cultures derived from these lines have previously been shown to display ALS pathological features (e.g. dipeptide repeat proteins (DPRs), RNA foci, protein mislocalization) and various cellular phenotypes (e.g. altered excitability, cell death) (Ruxandra Dafinca et al., 2016; Guo et al., 2017a; Naujock et al., 2016; Ramic et al., 2021; Sareen et al., 2013). All iPSC lines were generated from fibroblasts and were subjected to karyotyping. Detailed information regarding the iPSC lines is shown in **Table S1**. Functional spinal lower MN were generated using a modified version of a previously published protocol (Du et al., 2015; Zelina et al., 2024) (**Figure 1A**). iPSCs were differentiated into MNs in six-well plates, with all iPSC lines (n=18) organized into two different batches each containing 9-10 lines (**Figure 1B**). One healthy control (HC) line was included in both batches (HC-1). Each batch contained a mix of HC, ALS and isogenic (iso) control lines (iCTRL). To assess variability during the differentiation process, the iPSC lines within each batch were subjected to two independent differentiations (i.e. two technical replicates). All lines were successfully differentiated into MNs within 25 days (DIV25), in line with previous studies (Dafinca et al., 2020; Ruxandra Dafinca et al., 2016; N. Egawa et al., 2012; Guo et al., 2017a; Naujock et al., 2016; Sareen et al., 2013). Cultures contained HB9- and ISL1-positive MN, in addition to expression of CHAT and SMI32/NFH, as shown by immunofluorescence analysis (**Figure 1C, S1A**). These markers were further confirmed by quantitative (q)RT-PCR showing *HB9*, *ISL1*, *CHAT* and *SMI32* expression at DIV17, DIV25 and DIV30 (**Figure 1D**). Markers of iPSCs (*NANOG*, *SOX2*) and neural progenitors (*PAX6*) were low, whereas several other (motor) neuron markers displayed increased expression during differentiation. No obvious differences in the expression of MN markers or cell viability were detected between HC and ALS cultures at DIV25, consistent with previous studies (Naohiro Egawa et al., 2012; Guo et al., 2017a) (**Figure 1D, S1B**).

**Figure 1:**
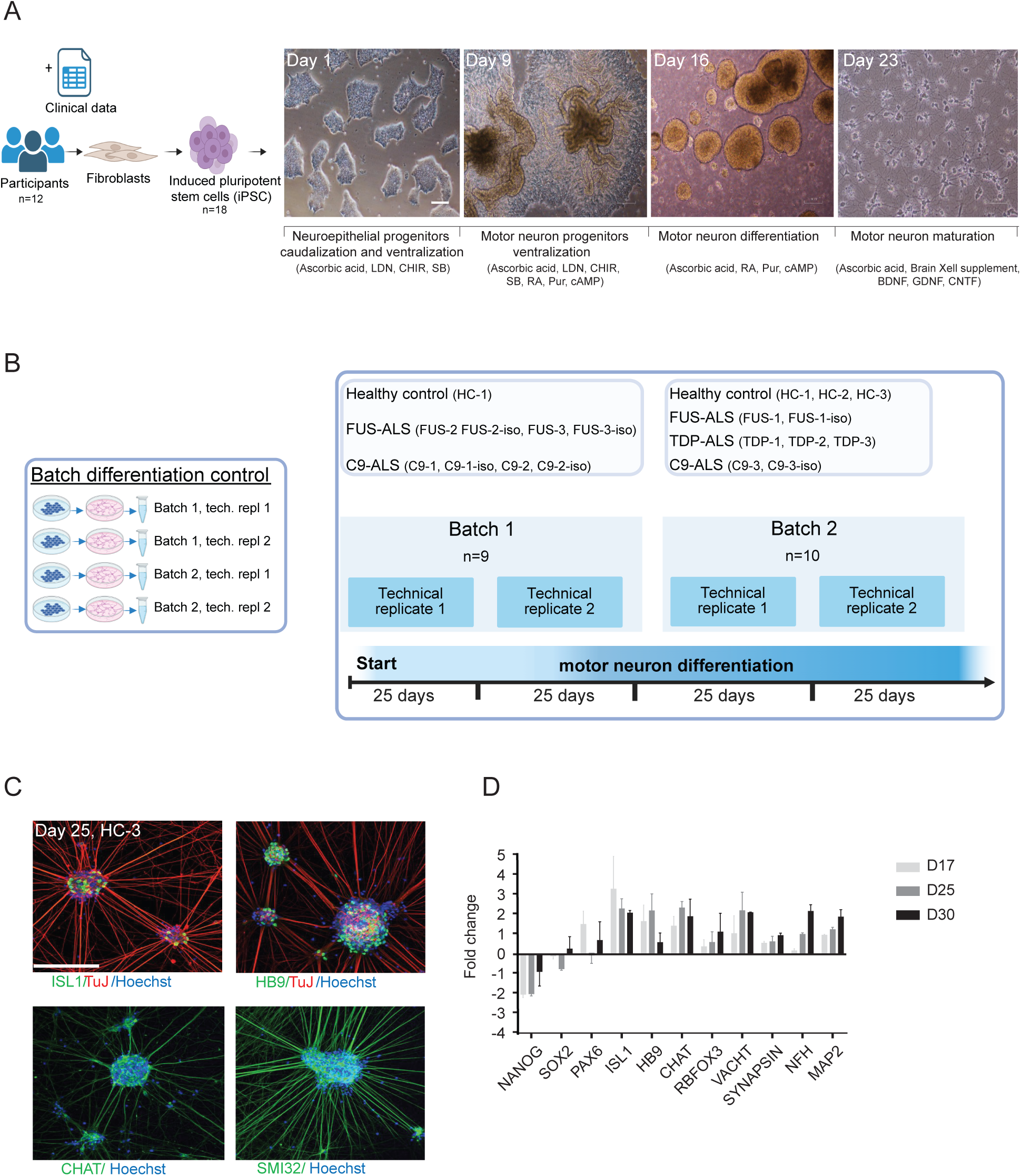
Differentiation and characterization of iPSC-derived motor neurons. **(A)** Schematic overview of the differentiation of iPSCs into motor neurons (MNs). Healthy control (HC), C9-ALS (carrying C9ORF72 repeat expansions), C9 isogenic control (C9-iso), FUS-ALS, FUS-iso, and TDP-ALS lines were differentiated into DIV25 MNs. **(B)** Schematic overview of sample distribution and batches. A total of 18 iPSC lines were divided into two batches, (Batch 1: n=9, Batch 2: n=10), with one HC line (HC-1) included in both batches. Each batch was differentiated two times (technical replicate) to control for technical variability. **(C)** Representative images of DIV25 MNs (HC-3) immunostained for 1) ISL1 (in green) and TuJ (in red), 2) HB9 (in green) and TuJ (in red), 3) CHAT (in green), and 4) SMI32 (in green). Hoechst is in blue. Scale bar, 250 µm. **(D)** mRNA expression levels of iPSC-derived MNs differentiated for 17, 25, and 30 days. The analysis includes expression of neural stem cell markers (*NANOG, SOX2, PAX6*), MN markers (*ISL1, HB9, CHAT, RBFOX3, VACHT, NFH*), and mature MN markers (*SYNAPSIN, MAP2*). Graph shows fold change values relative to gene expression in iPSCs, comparing two independent differentiations of iPSC-derived MNs (HC-3; n=2). Values > 1 indicate upregulation in MNs whereas values < 1 indicate downregulation relative to expression in iPSCs.

To resolve the proteome changes caused by different ALS-associated mutations in iPSC-derived MNs, high-resolution tandem mass spectrometry (LC-MS/MS) analysis was performed on the different samples outlined above. Up to 6209 proteins were identified with a false discovery rate (FDR) of 1%, with data demonstrating normal distribution across samples and replicates (**Figure S2A, S2B**). The different number of proteins identified per sample, ranging from 2833 to 5478 proteins, is likely due to variability in sample preparation. We removed 2 samples (C9-1 B1R1, C9-1-iso B1R1) due to insufficient protein identification by mass spectrometry. Annotation of the proteins revealed a wide variety of protein classes, including enzymes, membrane proteins, cytoskeletal proteins and gene-specific transcriptional regulators (**Figure S2C**). iPSC markers such as SOX2 and OCT4 were not detected, but samples showed strong expression of neuronal markers (e.g. GAP43, NEFL, MAP2, NCAM1). Further, FUS and TDP-43 were consistently expressed, whereas C9ORF72 was not detected, possibly due to low abundance (**Figure S2D**).

To evaluate overall protein expression patterns across lines and batches, principal component analysis (PCA) was performed. The first principal component (PC1) accounted for 11.8% of variation and PC2 for 7.1% **(Figure 2A**). Based on overall protein expression no clear separation was detected between ALS and control samples, nor between samples with different genotypes. A heatmap of the top 30 most variable proteins identified by PC1, revealed enrichment of ribosomal and RNA-binding proteins (**Figure 2B**). To further assess variation between batches and replicates, Pearson correlations were plotted for the HC-1 line, which was included in all batches and replicates. The correlation values ranged from 0.69 ≤ r ≤ 1, indicating that batch-to-batch variation is largely driven by biological variation reflecting individual-specific factors (**Figure 2C**). These observations align with findings from previous, large-scale studies of iPSC-derived MNs from ALS patients (Workman et al., 2023). Collectively, these results highlight the robustness of the differentiation protocol and reveal inter-individual variability in the proteomic landscape of iPSC-derived MNs.

**Figure 2:**
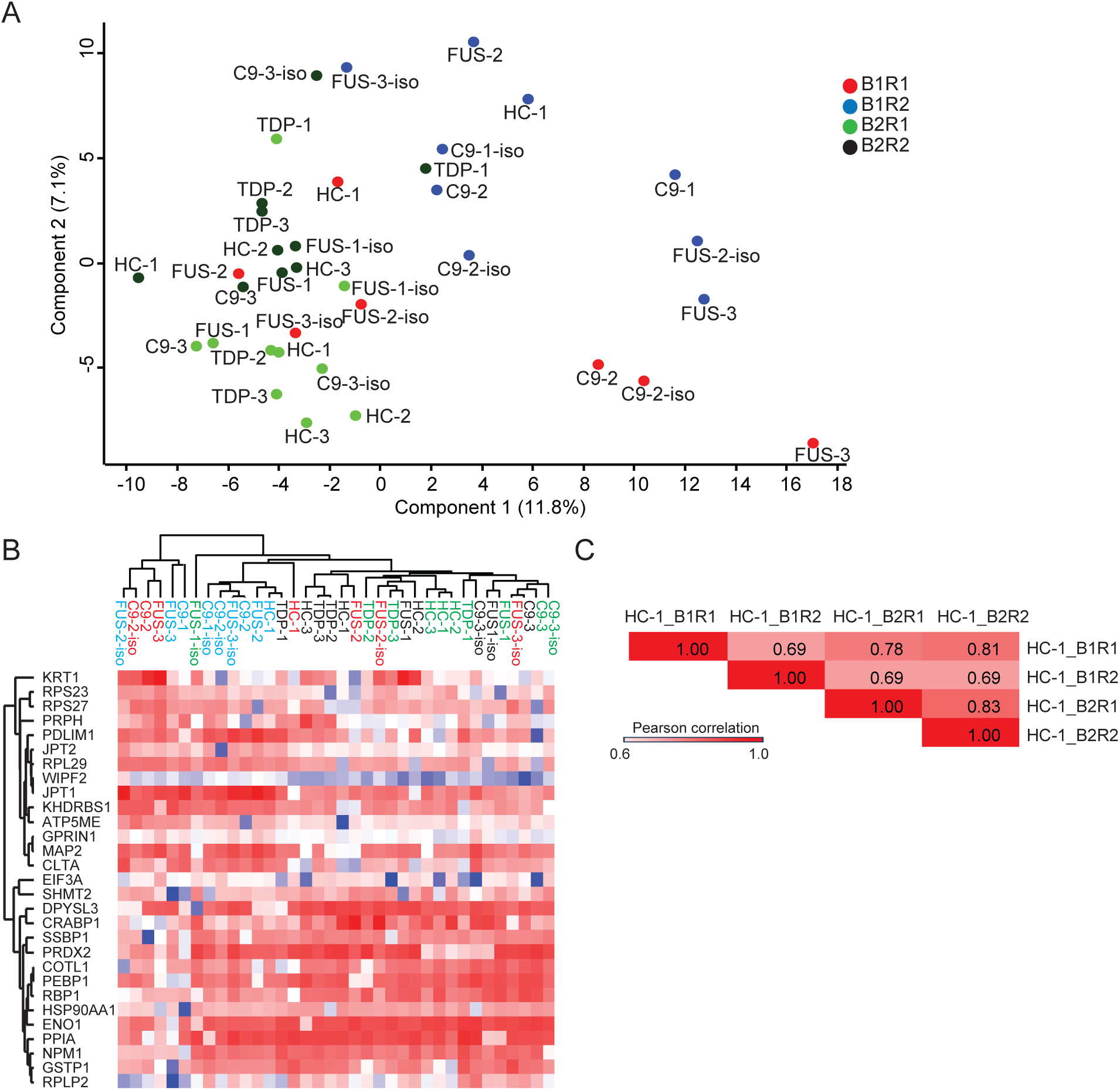
Proteomic analysis of iPSC-derived motor neurons carrying ALS mutations. **(A)** Principal component analysis (PCA) of the proteomes of DIV25 motor neuron (MN) samples, color-coded by batch (B) and replicate (R). **(B)** Heatmap showing the top 30 most variable proteins identified by PC1. **(C)** Pearson correlation analysis of the differentiation of healthy control iPSC line HC-1 across batches and replicates.

### ALS mutation-specific proteomic signatures in iPSC-derived MN

To determine ALS mutation-specific proteomic signatures, protein expression in C9-ALS, FUS-ALS or TDP-ALS samples was compared to HC. Differentially expressed proteins were defined using a *P*-value ≤ 0.1 and a fold change ≥ 1.3, selected based on data distribution to ensure a balance between statistical significance and biological relevance. This analysis revealed 75 upregulated and 199 downregulated proteins in C9-ALS MNs (**Figure 3A**), 62 upregulated and 136 downregulated proteins in FUS-ALS MNs (**Figure 3B**) and 74 upregulated and 66 downregulated proteins in TDP-ALS MNs (**Figure 3C**, **Table S2**). Gene Ontology (GO) enrichment analysis and KEGG pathway analysis were conducted to clarify the biological processes involved in each mutation genotype. For C9-ALS, GO analysis revealed upregulation of cellular response to nerve growth factor and vesicle-mediated transport, while mRNA splicing and metabolic processes were downregulated. KEGG pathways analysis confirmed the upregulation of synaptic vesicle cycle and downregulation of pathways associated to ALS (**Figure S3**). In FUS-ALS, several biological processes related to endoplasmic reticulum (ER) organization were both up- and downregulated. Additionally, pathways related to axon guidance and cGMP-PKG signaling were upregulated, while fatty acid degradation and ribosome-related pathways were downregulated. For TDP-ALS, upregulated GO terms included plasma membrane repair and cytoskeleton organization, with downregulation in MN axon guidance and fatty acid oxidation. KEGG pathway analysis related to endocytosis and oxidative phosphorylation were upregulated and several metabolism-related pathways were downregulated.

**Figure 3:**
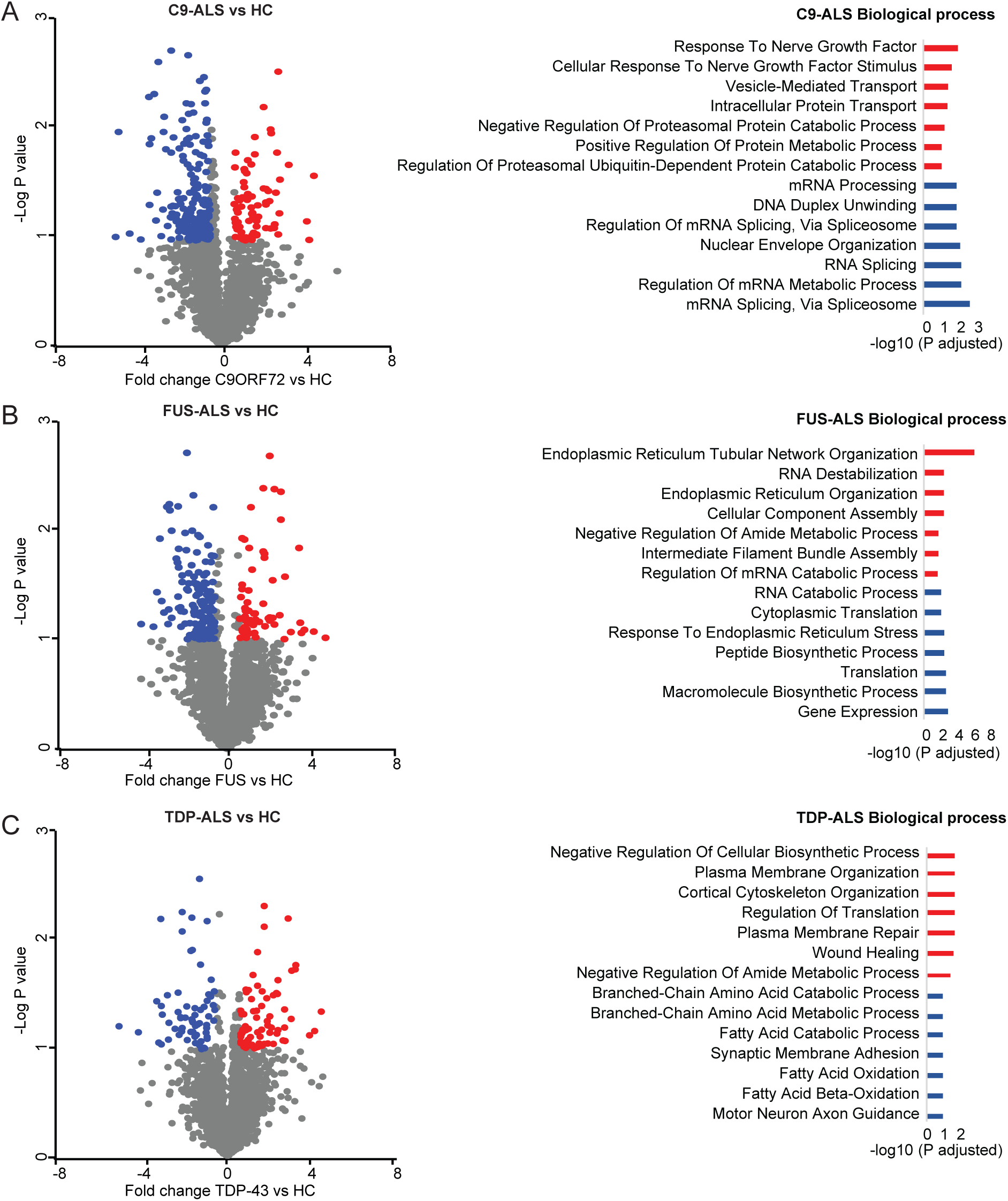
ALS mutation-specific proteomic changes in iPSC-derived motor neurons. Volcano plots showing differentially expressed proteins in C9-ALS (**A**), FUS-ALS (**B**) and TDP-ALS (**C**) motor neurons (MNs) at DIV25 as compared to healthy control (HC). The x-axis shows log2-fold change and y-axis -log10 (*P*-value). Proteins were considered significant based on a *P*-value cutoff of ≤ 0.1 and a fold change ≥ 1.3. Downregulated proteins are indicated in blue and upregulated proteins in red. GO analysis was performed to assess the biological functions of significantly altered proteins and a selection of GO terms (Biological process) is shown.

To further assess ALS mutation-specific changes, we compared the C9-ALS and FUS-ALS proteomes to that of isogenic controls (iCTRL), which were generated from these lines. This analysis revealed 48 upregulated and 99 downregulated proteins in C9-ALS and 95 upregulated and 35 downregulated proteins in FUS-ALS (**Figure 4A, B**). For C9-ALS, GO analysis revealed upregulation of oxidative phosphorylation and (mitochondrial) ATP synthesis, while ER stress-related processes were downregulated. In FUS-ALS, biological processes associated with cytoplasmic translation and gene expression were upregulated, whereas transmembrane transport activity and axonogenesis were downregulated. Interestingly, several proteins were significantly altered in comparisons to both HC or iCTRL samples (**Figure 4C**). Proteins exclusively altered in C9-ALS were involved in Golgi vesicle transport, protein localization to ER, and the regulation of neurogenesis. In FUS-ALS, such differentially expressed proteins were associated with intermediate filament bundle assembly, ER organization and RNA degradation. Overall, this analysis provides valuable insights into the molecular mechanisms underlying ALS related to *C9ORF72, FUS, and TDP-43* mutations, highlighting potential pathways for targeted therapeutic strategies.

**Figure 4:**
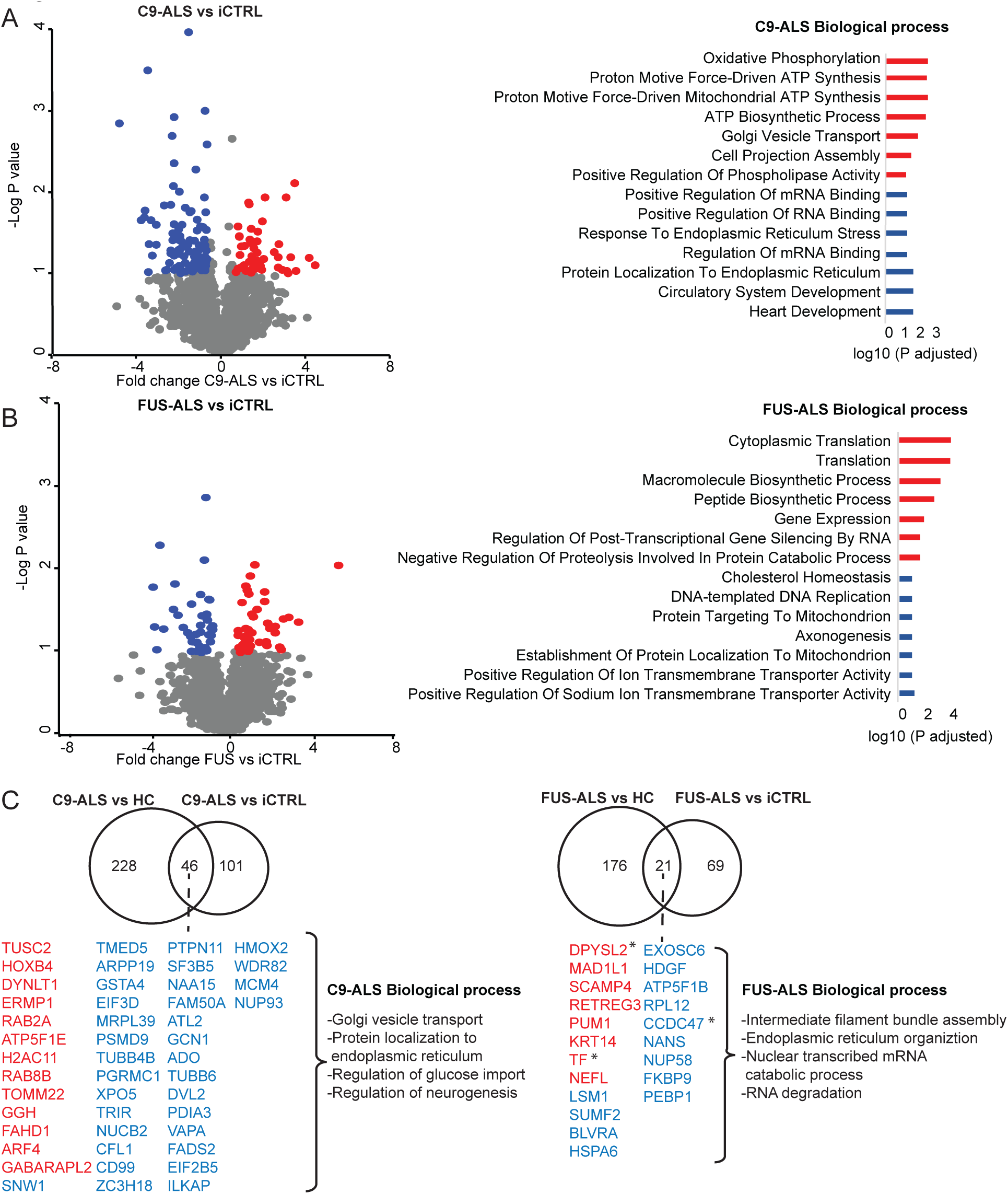
Proteomic changes in iPSC-derived motor neurons caused by C9ORF72 or FUS mutations. **(A)** Volcano plot showing differentially expressed proteins in C9-ALS motor neurons (MNs) compared to their isogenic controls (iCTRL) lacking the repeat expansion. **(B)** Volcano plot showing differentially expressed proteins in FUS-ALS MNs relative to their iCTRL. GO analysis were performed for significantly altered proteins (*P* ≤ 0.1, fold change ≥ 1.3). **(C)** Venn diagram illustrating shared dysregulated proteins in C9-ALS and FUS-ALS MNs, compared to healthy control (HC) or matching iCTRL. Downregulated proteins are indicated in blue and upregulated proteins in red. Asterisks indicate proteins regulated in the opposite direction in FUS-ALS versus iCTRL. Biological functions of these overlapping proteins are indicated.

### Common proteomic changes in ALS motor neurons

Following the characterization of mutation-specific proteome signatures in ALS, we sought to identify common proteomic alterations across these mutations. To achieve this, we combined data from C9-ALS, FUS-ALS, and TDP-ALS MNs and performed comparative analyses against HC samples. Proteins were considered significantly regulated based on a threshold of *P*-value ≤ 0.1 and a fold change ≥ 1.3. This integrative analysis identified 100 upregulated proteins and 64 downregulated proteins (**Figure 5A, Table S3**). GO analysis revealed upregulation of biological processes related to axonal transport and apoptosis, while oxidation-related processes were downregulated. Additionally, KEGG pathway analysis highlighted the upregulation of pathways involved in endocytosis and neurodegeneration, whereas pathways related to fatty acid degradation were downregulated. Notably, NEFL (neurofilament light chain), a critical biomarker in ALS pathology, was significantly upregulated in all conditions. Several of these dysregulated processes were also observed in some of the mutation-specific analyses. Analysis of the top 10 most significantly up- and downregulated proteins identified proteins implicated in apoptosis, cell adhesion and stem cell differentiation (**Figure 5B**). In conclusion, our integrative proteomic analysis highlights key shared molecular signatures across distinct ALS-associated mutations, offering insights into common pathogenic mechanisms at the protein level.

**Figure 5:**
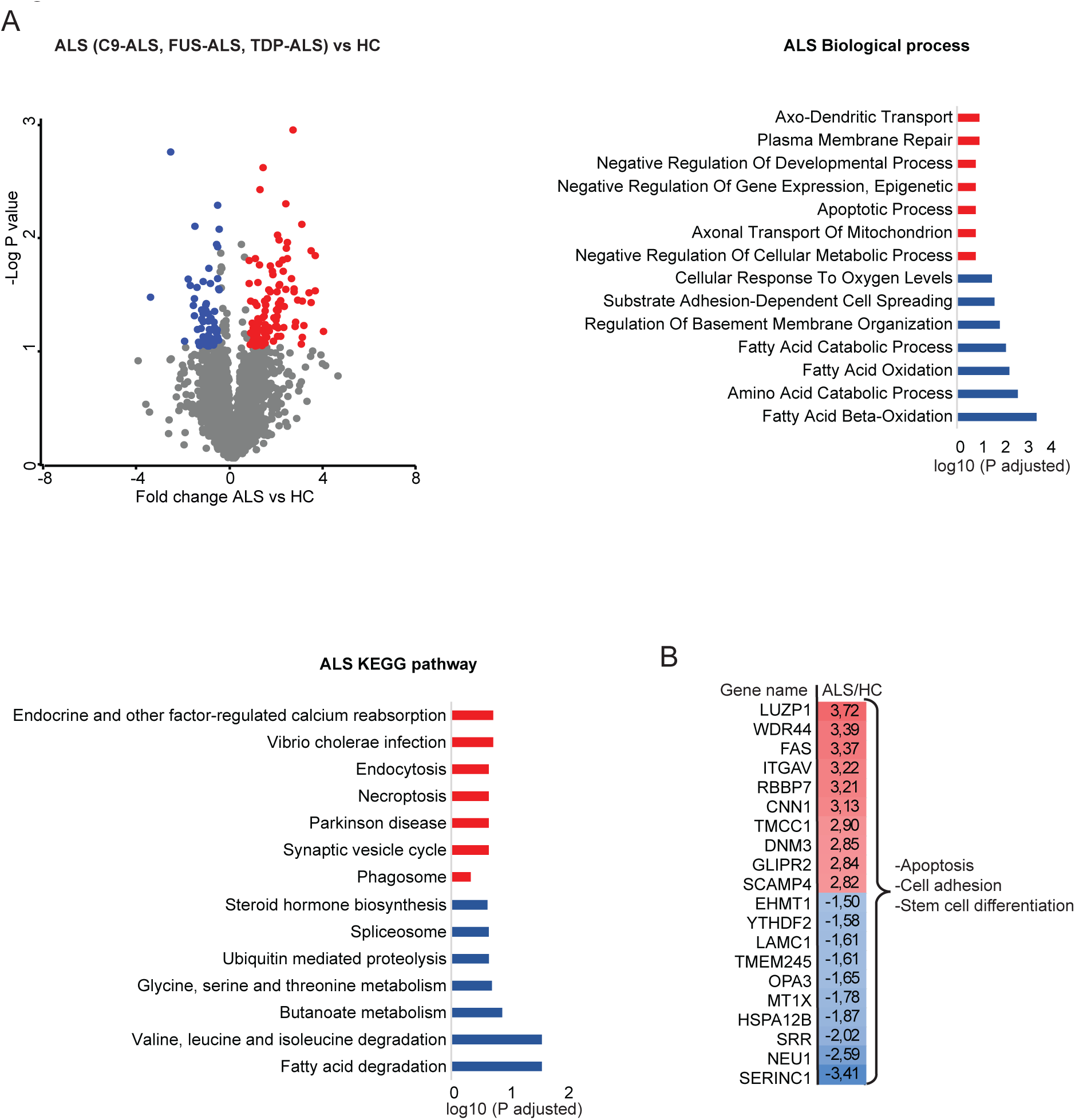
Common proteomic changes in ALS iPSC-derived motor neurons. **(A)** Volcano plot demonstrating differentially regulated proteins in ALS (C9-ALS, FUS-ALS and TDP-ALS) compared to healthy control (HC) motor neurons (MNs). The x-axis shows log2-fold change and the y-axis the -log10 (*P*-value). Proteins with a *P*-value cut-off of ≤ 0.1 and a fold change ≥ 1.3 were considered significant. Downregulated proteins are shown in blue and upregulated proteins in red. GO analysis and KEGG pathway analysis were performed for the significant proteins (Biological processes). **(B)** Heatmap showing the top ten most upregulated and downregulated proteins in ALS. Biological processes linked to these proteins are shown at the right.

### MSC-EVs can reverse ALS phenotypes in vitro

Differential protein expression patterns can be exploited to dissect disease pathways and identify therapeutic targets. In addition, proteomic signatures can be used to assess the effect of therapeutic interventions at the protein/pathway level. Although several studies have reported beneficial effects of MSCs in ALS rodent models (Lewis & Suzuki, 2014; Uccelli et al., 2012), their effect in human *in vitro* models of this disease is unknown. Further, the therapeutic effects of these cells are likely primarily mediated through their secretome. For example, by proteins present in EVs released by MSCs (Varderidou-Minasian & Lorenowicz, 2020). Interestingly, MSC-EVs are known to influence neurite growth and branching (Lopez-Verrilli et al., 2016) and several ALS genetic defects cause altered neurite growth *in vitro*. For example, cultured FUS-ALS MNs display shorter neurites (Stoklund Dittlau et al., 2021). Therefore, to establish whether MSC-EVs can reverse ALS-associated cellular phenotypes, their effect on a neurite growth defect observed in FUS-ALS MN cultures was tested.

MSCs were derived from bone marrow aspirated from third-party healthy donor, approved by the Dutch Central Committee on Research Involving Human Subjects (CCMO, Biobanking for MSC, NL41015.041.12) and EVs were isolated using differential ultracentrifugation steps (Vonk et al., 2018). Western blot analysis confirmed expression of the exosomal markers CD9 and CD63, and absence of calnexin, an integral protein of the ER (**Figure 6A**). Nanosight Particle Tracking analysis (NTA) indicated that the nominal size of MSC-EVs was approximately 142 nm with a concentration of 6.4×10^6^ particles per milliliter (**Figure S4A**). This is in line with our previous work using electron microscopy and NTA (Vonk et al., 2018). To assess the effect of MSC-EVs on the reduced neurite growth displayed by FUS-ALS MNs, FUS-ALS and iCTRL cultures (FUS-3, FUS-3-iso) were exposed to MSC-EVs (EVs derived from 4×10^6^ MSCs per well of a 24 well plate), in line with previous data showing therapeutic efficacy (Vonk et al., 2018). On both days *in vitro* DIV25 (prior treatment) and DIV27 (after two days of treatment), FUS-ALS MN cultures displayed reduced neurite length as compared to iCTRL in the absence of MSC-EV treatment (**Figure 6B**). Two-day treatment with MSC-EVs did not significantly alter neurite growth in iCTRL conditions but reversed the neurite length reduction observed in FUS-ALS MN cultures. Together, these experiments show that MSC-EVs can reverse ALS-associated phenotypes in human *in vitro* MN models.

**Figure 6:**
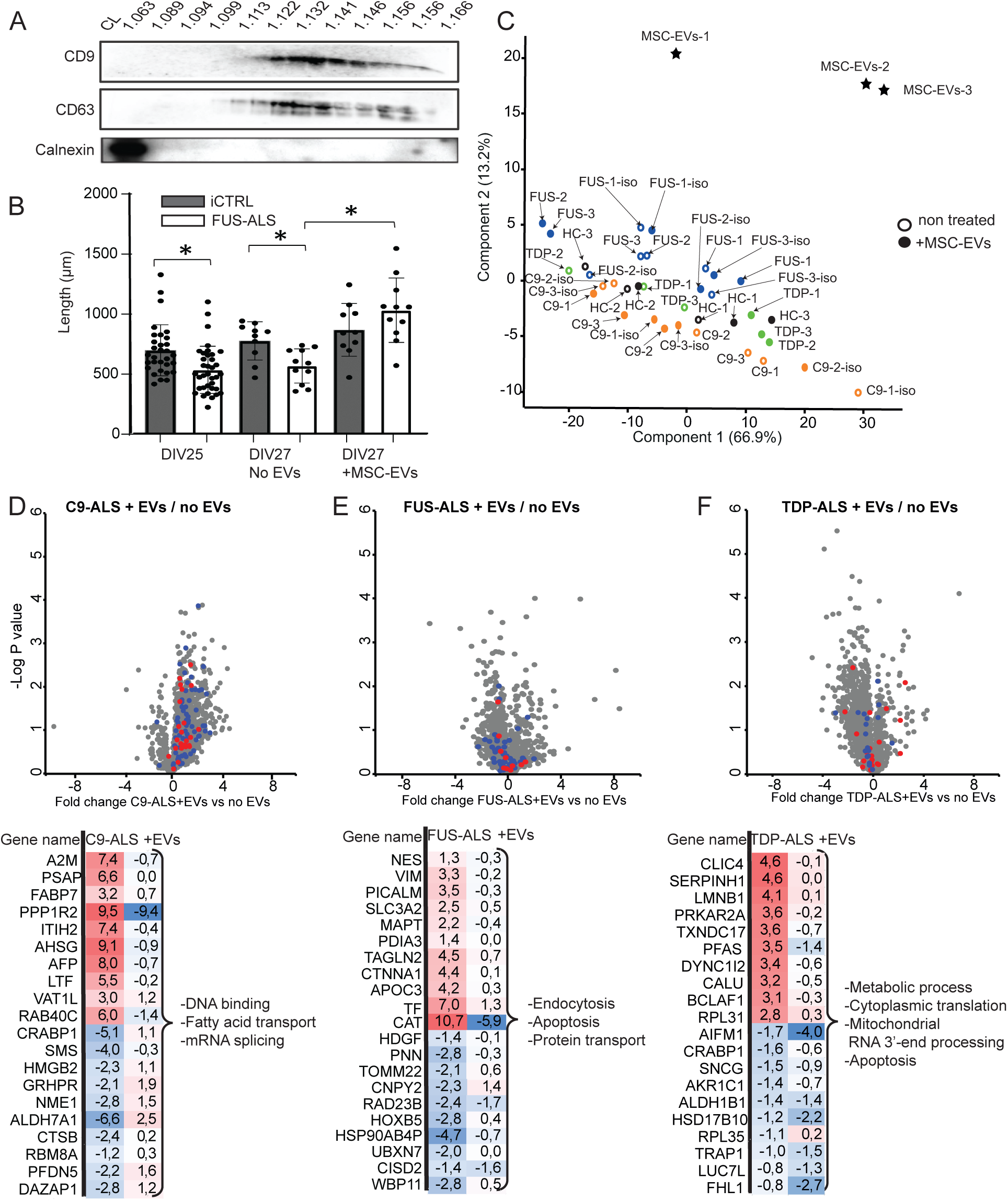
Treatment of iPSC-derived motor neurons with MSC-EVs. **(A)** MSC-EVs were fractionated using a sucrose density gradient (1.063–1.166 M), and sucrose density was confirmed using a refractometer. Twelve individual fractions were collected, along with cell lysate (CL) as a control. Western blot analysis detected the exosomal markers CD63 and CD9 in specific fractions, while the absence of the endoplasmic reticulum protein Calnexin confirmed minimal contamination by non-EV material. **(B)** Quantitative analysis of neurite length in FUS-ALS and iCTRL motor neurons (MNs) prior to and following two days of MSC-EV treatment. Data represent the mean ± SEM from three independent FUS-ALS lines (FUS-1, FUS-2, FUS-3) and their respective isogenic controls (FUS-1-iso, FUS-2-iso, FUS-3-iso) (N = 3). Statistical significance was determined using an unpaired two-tailed Student’s *t*-test. **(C)** Principal component analysis (PCA) of the proteomes of MSC-EVs (star), non-treated iPSC-derived MN samples (open circle) and MSC-EV-treated (closed circle) MN samples. Samples include ALS lines, healthy control (HC) and iCTRL **(D)** Volcano plot showing differentially expressed proteins in C9-ALS MNs treated with MSC-EVs compared to untreated MNs. Significantly regulated proteins in C9-ALS relative to HC are highlighted in red (upregulated) and blue (downregulated). **(E)** Volcano plot showing differentially expressed proteins in FUS-ALS MNs treated with MSC-EVs compared to untreated MNs. Significantly regulated proteins in FUS-ALS relative to HC are highlighted in red (upregulated) and blue (downregulated). **(F)** Volcano plot showing differentially expressed proteins in TDP-ALS MNs treated with MSC-EVs compared to untreated MNs. Significantly regulated proteins in TDP-ALS relative to HC are highlighted in red (upregulated) and blue (downregulated). Below each volcano plot the top ten most upregulated and downregulated proteins are shown together with the expression levels of these proteins after MSC-EV treatment. The Biological processes affected by MSC-EVs are shown at the right.

To begin to understand how MSC-EVs may affect ALS iPSC-derived MNs at the protein level, lysates from 3 independent MSC-EV isolations (100 µg) were subjected to mass spectrometry. A total of 1,967 proteins were identified with 1% FDR, including the exosomal markers CD63, CD81, and CD9 (calnexin was not detected). GO analysis of the MSC-EV proteome revealed enrichment in biological processes related to cellular localization, vesicle-mediated transport, and exocytosis (**Figure S4B**). The PCA plot further revealed a distinct protein composition in MSC-EVs compared to MNs (**Figure 6C**). Specifically, the MSC-EV proteome was enriched for defense immunity and cytoskeletal protein classes, while it contained fewer gene-specific transcriptional regulators (**Figure S4C**). Additionally, storage proteins such as ferritin were identified in MSC-EVs but absent in neuronal samples, suggesting differences in metabolic requirements between MSCs and neuronal cells. The top 25 most abundant MSC-EV proteins included albumin (ALB), thrombospondin-1 (THBS1), alpha-2-macroglobulin (A2M), fibronectin (FN1), and complement C3 (C3), which are known to be involved in immune modulation, extracellular matrix remodeling, and tissue repair (**Figure S4D**).

### Proteomic signature after MSC-EV treatment of MNs carrying specific ALS mutations

To explore the molecular mechanisms underlying the rescue of neurite growth phenotypes observed in FUS-ALS MNs by MSC-EVs, we performed proteomic analysis of MNs to examine changes in protein expression induced by MSC-EV treatment in MNs carrying three different ALS mutations. We focused on the proteins that were significantly up- or downregulated in ALS samples from each mutation (C9-ALS, FUS-ALS, TDP-ALS) and visualized these changes in volcano plots (**Figure 6D-F, Table S4**). Interestingly, levels of most significantly dysregulated proteins in ALS were either normalized or shifted toward the opposite direction after MSC-EV treatment. This suggests that MSC-EVs may help restore normal cellular function by modifying the expression of key proteins involved in ALS pathogenesis. This was confirmed by analysis of the top 10 most significantly up- and downregulated proteins for each mutation compared to HC and their normalized expression following MSC-EV treatment. For C9-ALS MNs, MSC-EV treatment predominantly affected DNA binding, fatty acid transport and mRNA splicing. For FUS-ALS, treatment had a major impact on endocytosis, apoptosis and protein transport. For TDP-ALS, MSC-EV exposure targeted metabolic processes and apoptotic pathways.

Next, common proteomic changes in ALS MNs were assessed following MSC-EV treatment (**Figure 7A, B**). As observed in mutation-specific analyses, treatment with MSC-EVs led to the normalization of the proteomic profile in ALS cells. Specifically, MSC-EV treatment targeted several key biological processes that were dysregulated across all ALS mutations, including oxidation, mRNA splicing and autophagy. These processes are commonly disrupted in ALS pathogenesis, and their restoration by MSC-EV treatment suggests a broad therapeutic potential for MSC-EVs in mitigating the molecular abnormalities shared across different ALS mutations.

**Figure 7:**
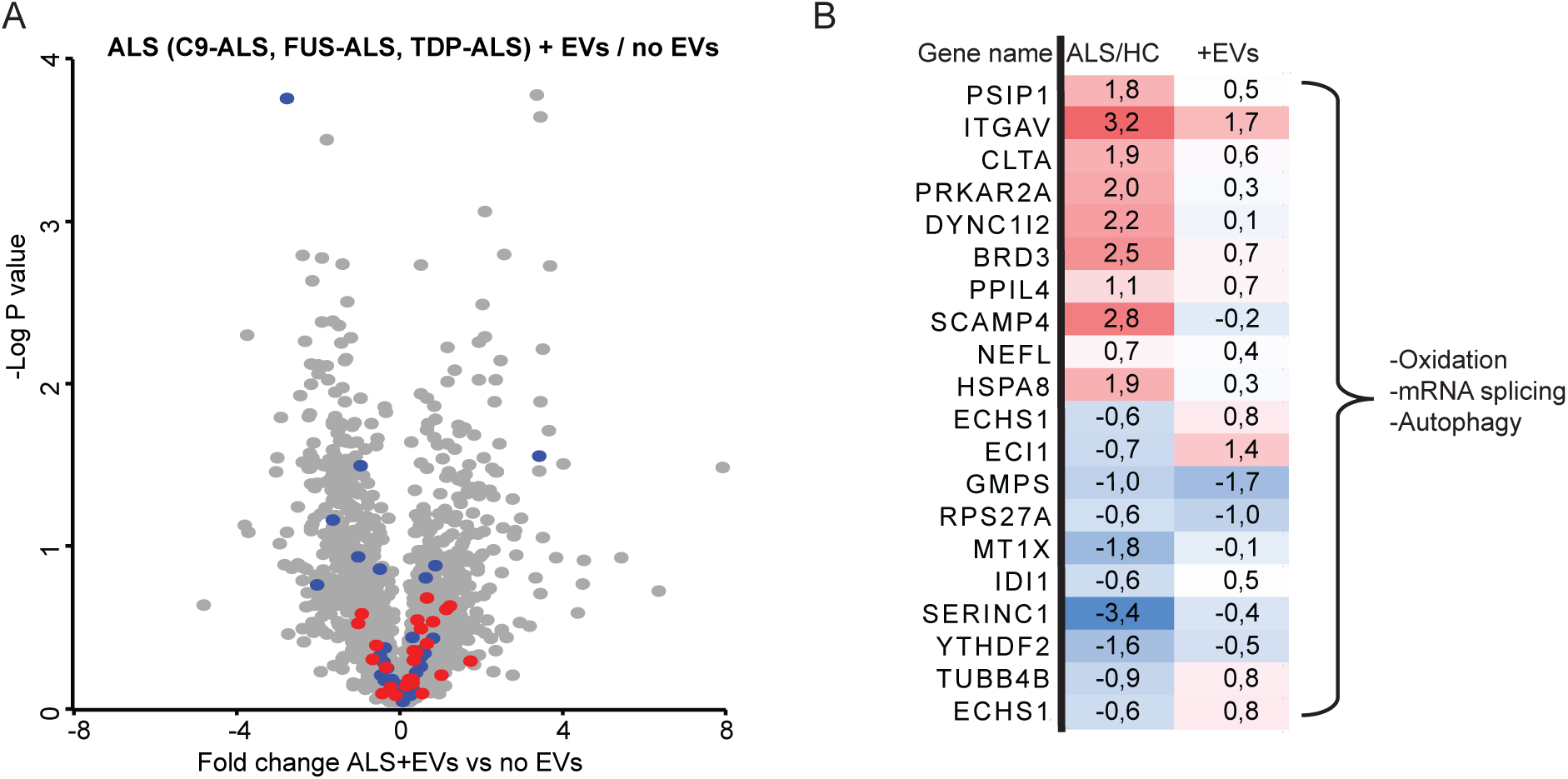
MSC-EV treatment normalizes ALS proteomic changes in iPSC-derived motor neurons. **(A)** Volcano plot showing differentially expressed proteins in ALS iPSC-derived motor neurons (MNs)(C9-ALS, FUS-ALS, and TDP-ALS) treated with MSC-EVs compared to non-treated MNs. Proteins that were significantly dysregulated in ALS compared to HC are shown in red (upregulated) or blue (downregulated). **(B)** The top ten most upregulated and downregulated proteins in ALS (C9-ALS, FUS-ALS, and TDP-ALS) are shown together with the expression levels of these proteins after MSC-EV treatment. The Biological processes affected by MSC-EVs are shown at the right.

## Discussion

iPSC-derived MNs are widely used to study ALS but our understanding of the disease-associated proteomic changes in these models is incomplete. These are, however, critical as they could serve as starting points for therapy development. Here, we conducted a comparative proteomic analysis of iPSC-derived MNs carrying ALS-linked mutations in *C9ORF72*, *TARDBP*, or *FUS*. This revealed both mutation-specific and shared proteomic signatures, unveiling common and divergent disease mechanisms. Further, it allowed us to show the ability of MSC-EVs to reverse ALS-associated proteomic changes in MNs carrying different gene mutations and restore functional defects in FUS-ALS neurons specifically. These findings unveil key proteomic signatures underlying ALS and identify MSC-EVs as a potential therapeutic approach for this disease.

### Mutation-specific and shared disease pathways in ALS motor neurons

Proteomic analysis of iPSC-derived MNs identified novel as well as previously reported mutation-specific changes. Several of the altered proteins belonged to pathways previously implicated in ALS, confirming the validity of our approach. In C9-ALS-specific MNs, we observed dysregulation of ER-Golgi vesicle transport, regulation of glucose import and neurogenesis. Disrupted ER-Golgi vesicle transport was previously reported in C9-ALS spinal cord tissues and iPSC-derived MNs (R. Dafinca et al., 2016; Prudencio et al., 2015). A reported role of C9ORF72 is to regulate membrane trafficking events, including ER-Golgi transport (Farg et al., 2014). Impaired ER-Golgi trafficking can disrupt the localization of proteins to the ER, leading to the accumulation of misfolded proteins, induction of ER stress, and activation of apoptotic pathways, all of which are upregulated in our C9-ALS MNs. Toxic DPRs produced from the C9ORF72 expansion may further contribute to these trafficking impairments and stress responses. In line, dysregulation of glucose import and downstream metabolism is disrupted at multiple levels—from brain region-specific uptake to intracellular mitochondrial metabolism (De Vocht et al., 2023; Wang et al., 2021). Impaired trafficking could also prevent proper membrane localization of glucose transporters such as GLUT3, limiting neuronal glucose uptake and have downstream consequences for processes such as neurogenesis. Changes associated with FUS-ALS included intermediate filament bundle assembly and RNA degradation. Abnormal assembly of intermediate filaments has been previously shown in the formation of axonal spheroids—focal swellings characterized by accumulation of neurofilaments—which are thought to disrupt axonal transport and contribute to neuronal dysfunction (Neumann et al., 2009). The disruption of axonal transport may underlie some of the morphological abnormalities frequently observed in FUS-ALS MNs. Interestingly, studies examining neurite architecture in FUS-ALS models have reported conflicting findings, with some showing increased axon branching and neurite complexity, while others report reduced neurite length and simplified arborization (Akiyama et al., 2019; Garone et al., 2021; Hawkins et al., 2022). These variations may reflect differences in disease stage, cellular context, or compensatory mechanisms. It is possible that early-stage FUS pathology leads to hyperbranching as a compensatory response to impaired transport, while progressive accumulation of cytoskeletal abnormalities eventually results in neurite shortening and degeneration. Furthermore, cytoplasmic aggregates of mutant FUS were shown to be the primary driver of dysregulated RNA splicing and RNA degradation, acting through both a loss of nuclear FUS function and a toxin gain-of-function (Rezvykh et al., 2023). Finally, TDP-ALS proteomic changes were linked to metabolic processes and downregulation of MN axon guidance. TDP-43 cytoplasmic aggregates were shown to localize to mitochondria and disrupt several respiratory chain complex proteins, resulting in energy deficiency and MN loss (Wang et al., 2016). Furthermore, the loss of nuclear TDP-43 resulted in increased inclusion of cryptic exons, leading to frameshifts and mRNA substrates for nonsense-mediated decay, a major pathway that regulates mRNA stability (Ling et al., 2015). However, the consequences of splicing factor, mis-localization and how this contributes to ALS pathogenesis require further investigation. These mutation-specific processes may be explained due to altered protein or RNA interactions. Previous interactome studies have shown that indeed, C9ORF72, TDP-43, or FUS display distinct protein and RNA interacting partners, that likely impair a range of distinct pathways underlying ALS pathology (Blokhuis et al., 2016).

Our data also show that mutations in these ALS-associated genes can trigger several common cellular pathways. We observed upregulation of apoptosis and axon-dendritic transport. It is important to note that dysfunction of the abovementioned mutation-specific processes can ultimately trigger general apoptotic processes. Despite well-documented impairments in axonal transport in ALS, transport of certain cargoes can be upregulated due to dysregulation of motor proteins. For example, mutations in the motor protein KIF5A disrupt autoinhibition, resulting in increased anterograde mitochondrial transport and contributing to ALS pathogenesis (Baron et al., 2022). In this study, we also report novel mechanisms that may contribute to ALS pathology, including the downregulation of fatty acid degradation—a topic that remains largely unexplored in ALS research. Notably, recent findings in C9-ALS also highlight this pathway as disrupted (Giblin et al., 2025). Given their high metabolic demands, MN rely heavily on mitochondrial β-oxidation of fatty acids to produce acetyl-CoA, which enters the TCA cycle to drive ATP synthesis. Impairment of this process may result in energy deficits that render MNs particularly vulnerable to cellular stress and degeneration. This mechanistic insight aligns with epidemiological evidence showing that higher dietary intake of fatty acids is associated with reduced ALS risk and improved survival after disease onset, supporting a potentially protective role for lipid metabolism in ALS progression (Veldink et al., 2007).

Several interesting candidates were identified among the top ten most significantly up- and downregulated proteins shared across the three ALS mutations. This included NEFL, which marks disrupted cytoskeletal integrity. NEFL accumulation is a recognized pathological hallmark of ALS and has been validated as a fluid biomarker for disease progression (Witzel et al., 2024; Wong et al., 2000). This strengthens the biological relevance of our proteomic findings and underscores the link between cytoskeletal dysfunction and MN degeneration in ALS. Moreover, its dual role as both a pathological marker and a quantifiable biomarker makes NEFL particularly valuable for monitoring disease activity and evaluating therapeutic responses in future studies. Together, our results indicate that the protein changes can serve as valuable starting point to better understand ALS pathogenesis and ultimately for the design of novel treatments.

### The therapeutic effect of MSC-EVs

Our data reveal several dysregulated cellular pathways, each likely contributing to the multifactorial pathogenesis of ALS. While most current experimental therapies target individual genes or pathways, our findings support a broader, multimodal therapeutic strategy. We demonstrate that MSC-EVs can have therapeutic effects in ALS MN cultures. Previous studies have shown that MSC-EVs improve motor performance in ALS mouse models (Bonafede et al., 2016; Lee et al., 2016). However, the underlying molecular mechanisms remained largely unknown. We exploited our proteomics data and approach to investigate the effect of MSC-EVs on iPSC-derived MNs from ALS patients with diverse genetic backgrounds. Our analysis revealed that MSC-EVs carry heterogeneous cargo, including enzymes, immune-related proteins, and transcriptional regulators, enabling them to simultaneously modulate multiple cellular pathways. This multimodal effect was supported by our proteomics data, which showed that MSC-EV treatment reversed both mutation-specific and commonly dysregulated protein changes in ALS MNs. For example, in FUS-ALS MNs, MSC-EV treatment normalized proteins associated with endocytic trafficking and apoptosis—key processes involved in cytoskeletal stability, axonal transport, and neuronal survival—which may underlie the observed rescue of neurite outgrowth defects. In C9-ALS MNs, MSC-EVs restored protein levels related to ER-to-Golgi vesicle transport, while in TDP-ALS MNs, MSC-EVs modulated proteins linked to metabolic processes and apoptosis. These findings are consistent with prior studies reporting that MSC-EVs exert neuroprotective effects through modulation of apoptosis, inflammation, and axonal growth (Bonafede et al., 2019; Giunti et al., 2021; Lopez-Verrilli et al., 2016). Importantly, we also treated HC and iCTRL MNs with MSC-EVs and did not observe a significant overall increase in the expression of the proteins that were altered in ALS MNs. This suggests that the uptake of MSC-EV cargo by healthy neurons does not result in a harmful accumulation of proteins. One possible explanation is that healthy cells possess efficient proteostasis and clearance mechanisms, such as the ubiquitin-proteasome system and autophagy, which allow them to selectively degrade or recycle excess proteins delivered by MSC-EVs. In contrast, ALS MNs, which are already under proteostatic stress, may benefit more from the exogenous support provided by MSC-EV proteins. These subtype-specific responses to MSC-EVs highlight the adaptability of MSC-EVs and underscore their potential as a precision-based, multimodal therapeutic agent capable of accommodating the heterogeneity of ALS. Beyond direct molecular effects, MSC-EVs may also exert indirect influences. For example, MSC-EVs may modulate the MN secretome, thereby altering the extracellular environment to promote the survival and function of neighboring MNs. Translation of MSC-EV therapy to ALS clinical application will require addressing several key challenges, including the standardization of EV production and dosing protocols, a deeper mechanistic understanding of their active components, development of effective delivery strategies to the central nervous system, and evaluation of long-term safety. Incorporation of MSC-EVs into combinatorial treatment strategies may enhance their impact and offer new avenues for disease modification.

In conclusion, our data reveal both mutation-specific and shared proteomic signatures linked to different ALS mutations. Some of these findings confirm pathways previously implicated ALS while others provide a valuable framework for dissecting novel disease mechanisms. In addition to being exploited for dissecting disease pathways or for therapy development, differential protein expression patterns can be used to assess the effect of therapeutic interventions at the protein/pathway level. In line with this, we show the ability of MSC-EVs to reverse proteomic changes in MNs with different ALS genetic backgrounds, in addition to being able to rescue axonal deficits in FUS-ALS. These findings highlight key molecular pathways in ALS at the protein level and support the potential of MSC-EVs as a possible therapeutic approach in ALS.

## STAR Methods

Detailed methods are provided in the STAR methods.

## Methods

### Cell culture

#### iPSC lines and culture

The iPSC lines used in this study, either generated at Utrecht Medical Center Utrecht or provided by others, were reprogrammed from skin fibroblasts. A detailed list of the iPSC lines used is provided in **Table S1.** All experimental procedures involving iPSCs were approved by the Medical Ethical Committee of the University Medical Center Utrecht. The C9ORF72 and HC iPSC lines have been previously characterized and reported (De Decker et al., 2024; van der Geest et al., 2024). iPSCs were cultured on Geltrex-coated dishes, passaged weekly using Trypsin-EDTA, and maintained in StemFlex^TM^ medium supplemented with ROCK-inhibitor Y-27632 for the first 24 hours after passage. Cells were frozen in fetal bovine serum (FBS) containing 10% dimethyl sulfoxide in cryotubes and cryopreserved in liquid nitrogen until further use. Cryopreserved cells were thawed at 37⁰C and washed with cold DMEM/F-12. iPSCs were used up to passage number 55. The pluripotency of all iPSC lines was confirmed by immunocytochemistry using the StemLight Pluripotency Antibody kit and the STEMdiff^TM^ Trilineage Differentiation Kit, as well as by quantitative reverse transcription PCR (qRT-PCR). Medium was replaced every other day and all lines were frequently tested for mycoplasma contamination using the MycoAlert kit.

#### Motor neuron differentiation

Spinal MN differentiation from iPSCs was performed according to a previously described protocol with some modifications (Du et al., 2015). Briefly, iPSCs were dissociated with 1mg/mL Dispase and plated on Geltrex-coated 6-well plates. Seeding density was optimized for each iPSC line depending on its proliferation rate. Basal medium was used throughout the differentiation process which consisted of a 1:1 mixture of DMEM/F12 and Neurobasal media, supplemented with 1x Sodium Pyruvate, 100 µM Non-Essential Amino Acids, 1x GlutaMAX, 1x Penicillin/Streptomycin, 1x N2 supplement, and 1x B27 supplement. Approximately 1-2 days after plating, medium was changed with basal medium supplemented with ascorbic acid (0.4 mM), LDN193189 (0.2 μM), CHIR99021 (3 mM), and SB431542 (10 μM,) for 6 days, with medium changes every other day. At DIV7, medium was changed to basal medium supplemented with ascorbic acid (0.4 mM), LDN193189 (0.2 μM), CHIR99021 (1 μM), SB431542 (10 μM), Retinoic Acid (RA) (200 nM), Purmorphamine (0.5 mM,) and cAMP (0.5 µM) for another 6 days, with daily medium changes. At DIV12, iPSCs were differentiated into MN progenitors, which were dissociated with Accutase and cultured in suspension in basal medium supplemented with ascorbic acid (0.4mM), cAMP (0.5 µM), retinoic acid (500 nM), and Purmorphamine (0.1 µM) for 6 days, with medium changes every other day. At DIV18, cells in suspension were washed in PBS and dissociated with Accutase for 30-45 min at 37°C into single cells and plated on coverslips coated with 20 µg/mL Poly-D-lysine (PDL) and 10 µg/mL Laminin. Adherent MNs were cultured in basal medium supplemented with ascorbic acid (0.4 mM), cAMP (1 µM), BDNF, GDNT and CNTF (10 ng/mL), and BrainXell supplement (diluted 1:1000). Medium was changed twice a week. Characterization and treatments were performed at DIV24-DIV25.

#### Mesenchymal stromal/stem cell culture

Human bone marrow samples were obtained from a non-HLA-matched healthy donor, with approval from the Dutch Central Committee on Research Involving Human Subjects (CCMO; Biobanking bone marrow for MSC expansion, NL41015.041.12). Bone marrow was isolated through density gradient centrifugation using Lymphoprep, and MSCs were isolated by plastic adherence, as previously described (Meng et al., 2016). MSCs were cultured in alpha-modified Eagle’s medium (α-MEM)) supplemented with 1% L-Glutamine, 100 U/mL penicillin, 100 µg/mL streptomycin, 5% human platelet lysate, and 10 U/mL heparin (Prins et al., 2009). Medium was changed every 3 days. Upon reaching 80% confluency, MSCs were washed with PBS, detached using 0.05% Trypsin-EDTA, and cultured up to passage 3 in 175 cm^2^ culture flasks.

### Extracellular vesicle isolation

EVs were isolated from MSC culture medium as described previously (Théry et al., 2006). Briefly, MSCs were cultured for 48 hours in medium which was depleted from EVs by overnight centrifugation at 100.000 x g. EVs were then isolated from conditioned medium through a series of centrifugation steps. First, cells were removed by two centrifugation steps at 200 x g for 10 minutes. Then, supernatant was centrifuged twice at 500 x g for 10 minutes, followed by 10.000 x g for 45 minutes. EVs were then pelleted by ultracentrifugation at 100.000 x g for 16 hours at 4 °C using SW32Ti rotor (Beckman). Finally, EVs were washed and pelleted at 100.000 x g overnight in PBS containing 0.5% BSA using a SW60 rotor (Beckman). EVs were stored in PBS with 0.5% BSA at -80°C until further use. MNs were treated with EVs derived from 4 x 10^6^ MSCs (or 3 x 10^8^ particles) per well in a 24 well plate for 2 days.

### Sucrose density gradient

EVs were resuspended in 250 μL PBS containing 2.5 M sucrose and transferred to a SW60 tube. The EV suspension was layered with 15 sequential 250 μL layers of 20 mM Tris-HCl (pH 7.4), each containing decreasing concentrations of sucrose, ranging from 2 M to 0.4 M. The gradient was then subjected to ultracentrifugation at 200.000 x g for 16 hours at 4°C. After centrifugation, 250 μL fractions were collected, and sucrose density was determined using a refractometer.

### Nanoparticle Tracking Analysis

The size distribution and quantification of isolated EVs was determined using Nanosight NS500 (Marvern, Worcestershire, UK) with the following settings: camera level: 13, slider shutter: 1232, slider gain: 219, detection: 5, autoblur mode for the analysis. Samples were diluted 100, 300, 500, 700, and 1000 times with PBS to reach an optimal concentration for instrument linearity. Data was processed using NTA Software v3.1.54 employing the finite-track-length adjusted algorithm with 10 nm bin widths. Size distribution and concentrations were calculated using build-in-software.

### RNA isolation, cDNA synthesis and qPCR

Cells were lysed using TRIzol reagent, and RNA was extracted using the RNeasy Mini Kit according to the manufacturer’s instructions, including on-column DNAse treatment. RNA concentration and purity were measured in a NanoDrop 2000 Spectrophotometer. Intron-spanning primers were designed with PrimerBLAST (NCBI) (**Table S5**). cDNA synthesis was performed using the SuperScript IV Reverse Transcriptase Kit, following the manufacturer’s instructions. qPCR reactions were performed with 2x FastStart Universal SYBR Green Master in a QuantStudio 6 Flex System qPCR cycler, using the standard PCR program. Each sample was analyzed in duplicate, and expression fold changes were calculated using the 2^−ΔΔCT^ method. Data were normalized to the geometric mean of the three reference genes: *GAPDH*, *BETA-ACTIN* and *TBPQ*. Graphical analysis was performed using GraphPad Prism.

### Immunocytochemistry

Cells were fixed in 4% paraformaldehyde in PBS for 20 min at 4°C and permeabilized with 0.1% Triton-X-100 in PBS for 5 min at room temperature (RT). Blocking was performed using 2% BSA, 0.1% Triton X-100 and 5% goat serum in PBS for 30 minutes at RT. Primary antibodies were diluted in blocking buffer and incubated overnight at 4°C. Following three washes with 0.1% Tween in PBS, slides were incubated with species-specific Alexa Fluor 488-, Alexa Fluor 568-, or Alexa Fluor 647-conjugated secondary antibodies (1:500) for 1 hour at RT. Nuclei were counterstained with DAPI for 10 minutes. Cells were mounted with FluorSave and imaged with confocal microscope.

### Westernblot

Cell pellets were lysed in RIPA buffer (25 mM Tris-HCl, pH 7.6, 150 mM NaCl, 1% NP40 and 1% NaDOC) containing 0.1% SDS, Complete Protease Inhibitor Cocktail, and 1x NuPAGE loading buffer, and incubated for 5 min at 95°C. Protein concentration was determined using the microBCA Protein Assay Kit following the manufacturer’s protocol. For SDS-PAGE, proteins were separated and transferred to a polyvinylidene difluoride (PVDF) membrane according to the manufacturer’s instructions. The membrane was blocked with 5% BSA for 1 hour at RT. After blocking, blots were incubated with primary antibodies in 5% BSA overnight at 4°C. The following antibodies were used: mouse anti-CD9 (1:1000), mouse anti-CD63 (1:1000) and rabbit anti-calnexin (1:1000). After washing, the blots incubated with appropriate peroxidase-conjugated secondary antibodies in 5% BSA for 1 hour at RT. Immunoreactivity was detected using chemiluminescence reagents, and signals were detected with the ImageQuant LAS 4000 Biomolecular Imager (GE Healthcare Life Sciences).

### Proteomics

#### Sample preparation

Samples were collected and lysed with sodium deoxycholate (SDS) lysis buffer containing 1% SDC, 10 mM tris(2-carboxyethyl)phosphine hydrochloride (TCEP), 40 mM chloroacetamide (CAA), and 100 mM TRIS, pH 8.0, supplemented with protease inhibitor cocktail. The lysate was sonicated using a Bioruptor (Diagenode) and centrifuged at 2.500 x g for 10 min at 4°C. A total of 20 µg protein per sample was digested overnight with Trypsin (1:50 protein-to-enzyme ratio) at 37°C. The digestion was quenched with 2% formic acid (FA). The resulting peptides were centrifuged at 20.000 x g for 10 minutes and vacuum-centrifuged to dryness. The digestion was acidified and desalted using C18 cartridges on the AssayMap BRAVO Platform (Agilent Technologies). Samples were dried and resuspended in 50 mM triethylammonium bicarbonate at a final concentration of 5 mg/mL prior to LC-MS analysis.

#### Mass spectrometry analysis

Mass spectrometry was performed in the Orbitrap Q-Exactive HF-X mass spectrometer (Thermo Scientific) coupled to reversed-phase nano-LC-MS/MS using Ultimate 3000 ultrahigh-performance liquid chromatography (UHPLC). Peptides were first separated on a 50 cm reversed-phase column (Agilent Poroshell EC-C18, 2.7 µm, 50 cm x 75 µm, in house packed) and eluted using a linear gradient with buffer A (0.1% FA) and buffer B (80% acetonitrile, 0.1% FA). Peptides were eluted over a 120-minute gradient from 13% to 44% buffer B at a flow rate of 300 nL/min. A column wash and re-equilibration step was followed with 55 minutes of total data acquisition time. The Q-Exactive HF-X mass spectrometer was operated in data-independent acquisition (DIA) mode, with a *m/z* range of 345-1650. Full-scan spectra were recorded at a resolution of 120.000 with an automatic gain control (AGC) target of 3 x 10^6^ with a maximum injection time of 100 ms. The normalized collision energy (NCE) was set to 27%, and the default charge state was set to 3.

#### Data processing

The protein sequence for *Homo Sapiens* (UP000005640) was downloaded from UniProt. Raw files were processed using DIA_NN (version 1.8.1) to generate spectral libraries via a deep-learning algorithm with the following parameters: Trypsin digestion with up to two missed cleavage sites, precursor m/z range of 300-1800, fragment ion m/z range of 200-1800, precursor charge range of 1-4, peptide length range 7-30 amino acids. N-terminal methionine excision and cysteine carbamidomethylation were enabled. Default settings were used including a false discovery rate (FDR) of 1% and match between runs (MBR) was enabled. The resulting *pg_matrix.tsv* file was used for further analysis.

#### Data visualization

Normalized protein and peptide abundances were extracted from Proteome Discoverer (PD2.2) and further analyzed using the open software PERSEUS environment for statistical and bioinformatics analysis and to generate the plots and figures. For several plots, GraphPad Prism was used. GO analysis was performed using Enrichr (Maayanlab.com). PANTHER pathway classification system (http://www.pantherdb.org/pathway/) was used for protein class analysis.

### Statistics

Data were normalized to the median and log2 transformed using PERSEUS software. Two samples (C9-1 B1R1, C9-1_iso B1R1) were excluded due to insufficient sample identification. Volcano plots were generated for the specified conditions, with proteins classified as up- or downregulated based on differentially expression criteria: a *P*-value ≤ 0.1 and a fold change ≥ 1.3. These thresholds were selected to balance statistical significance with biological relevance based on the data distribution. To study ALS mutation-specific alterations, C9-ALS, FUS-ALS or TDP-ALS samples were categorized in one group and compared to HC or iCTRL samples. For general ALS-related alterations, C9-ALS, FUS-ALS and TDP-ALS samples were categorized into a single group and compared to HC samples.

For GraphPad Prism analysis, a minimum of two independent differentiations were used for each biological condition. The results are presented as group mean with standard error of the mean (SEM), and statistical significance was determined using a two-sided t-test. The number of samples, replicates, and experiments are detailed in the Figure legends.

## Resource availability

### Lead contact

Jeroen Pasterkamp

### Material availability

This study did not generate new unique reagents.

### Data and code availability

All mass spectrometry proteomics data have been deposited to the ProteomeXchange Consortium via the PRIDE partner repository with the data set identifier PXD034824.

Username: Password: IDoEOWRj

## Acknowledgements

This work was supported by Stichting ALS Nederland (TOTALS, MUSALS, ATAXALS, GoALS), ALS CURE project, the INTEGRALS and MAXOMOD consortia (E-Rare-3, the ERANet for Research on Rare Diseases), TRIAGE (to R.J.P.) and NWO-XS (to S.V-M). We also would like to thank Kelly Stecker and Francine Rodrigues for their support and guidance with the mass spectrometry experiments.

## Author contributions

S.V-M designed and performed experiments and performed manuscript writing under supervision of R.J.P. with input from other authors. S.P-V. and A.T. performed qPCRs and immunostaining. C.J. performed immunostaining. M.A. supervised mass spectrometry proteomics. M.L. supervised the MSC-EV experiments. R.J.P. coordinated the study, acquired funding and wrote the manuscript with S.V-M.

## Supplemental Figure legends

**Figure S1: Characterization of iPSC-derived motor neuron cultures.**

Representative images of DIV24 iPSC-derived motor neurons (MNs) from different control and ALS iPSC lines immunostained for iPSC markers (NANOG) and MN markers (ISL1, TuJ, HB9, CHAT and SMI32). Hoechst is in blue. Scale bar is 250 µm.

**(A)** Cell viability of MNs with different genetic backgrounds was assessed using Trypan Blue. Each data point represents an independent differentiation. HC, healthy control; iso, isogenic control.

**Figure S2: Proteomic analysis of iPSC-derived motor neuron cultures.**

**(A)** Graph showing the number of identified proteins per sample.

**(B)** Graph showing Log2 transformed data per sample.

**(C)** Pie chart displaying the distribution of protein classes, based on PANTHER classification, for all quantified proteins (in all samples).

**(D)** Heatmap showing expression of marker proteins for iPSCs, progenitors, neurons, motor neurons, non-neuronal cells in addition to a few ALS-associated proteins. HC, healthy control; iso, isogenic control.

**Figure S3: KEGG pathway analysis of proteomic changes in iPSC-derived motor neurons carrying specific ALS mutations.**

KEGG pathway analysis was performed to identify altered molecular and cellular pathways based on significantly altered proteins per mutation (C9-ALS, FUS-ALS, and TDP-ALS) compared to healthy control (HC).

**Figure S4: Proteomic characterization of MSC-EVs.**

**(A)** Nanosight Particle Tracking Analysis (five independent measurements) of MSC-EVs showing that the nominal size of MSC-EVs is 142 nm. The red color indicates SEM.

**(B)** GO analysis based on the MSC-EV proteome (Biological process).

**(C)** Pie chart illustrating the distribution of protein classes among all quantified MSC-EV proteins, based on PANTHER classification. Data represent proteins identified from three independent MSC-EV isolations (n = 3).

**(D)** Top 25 most abundant proteins in MSC-EVs.

