## Supplementary figures and images for "Proteomic signature and mesenchymal stromal/stem cell-derived extracellular vesicle treatment of iPSC motor neurons from different ALS patient groups"

### Figure S1

Supplementary Figure 1

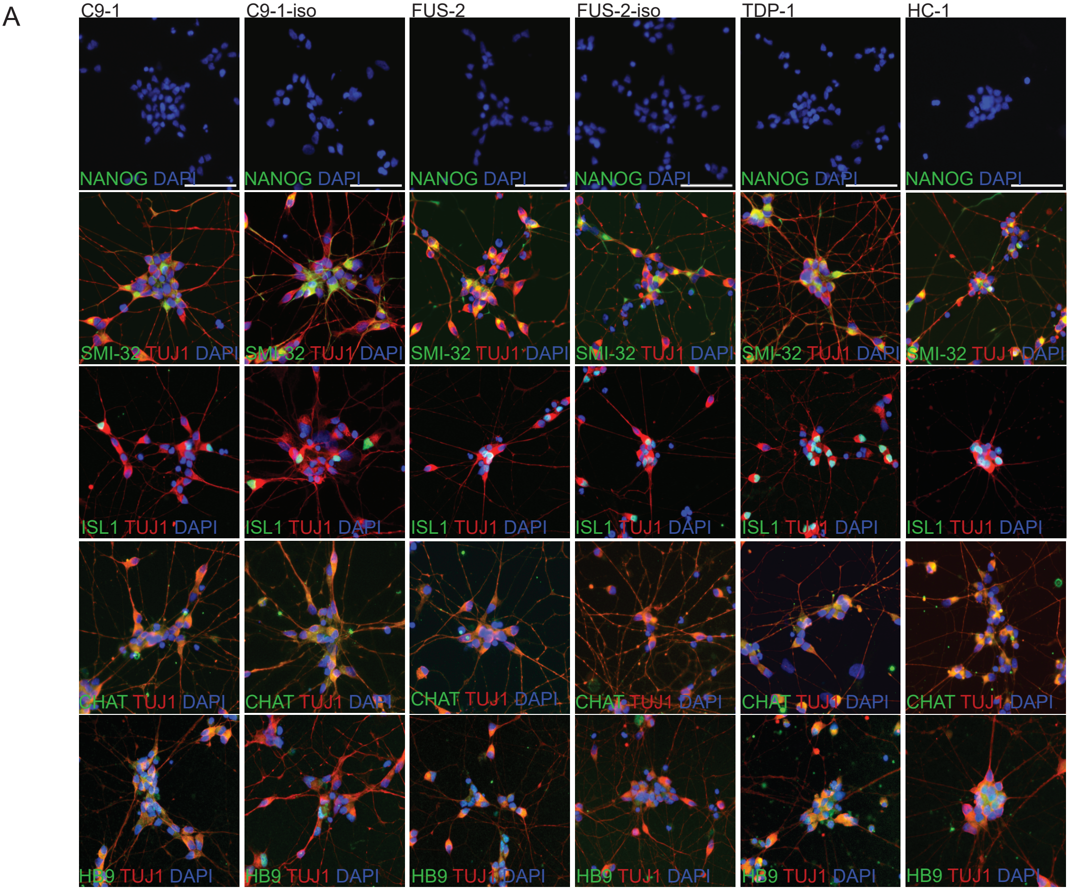


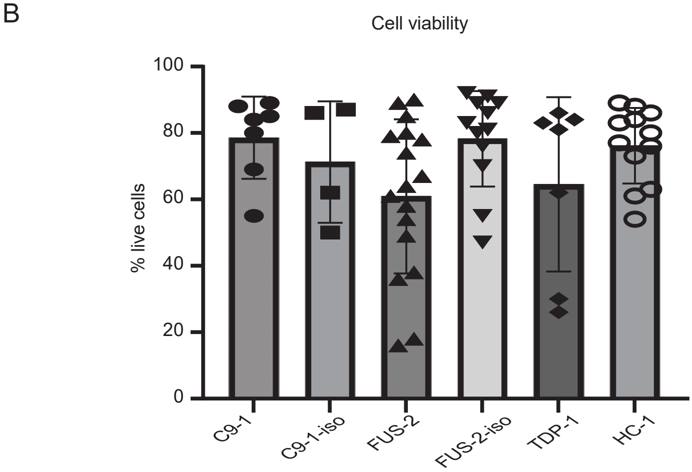

### Figure S2

Supplementary Figure 2

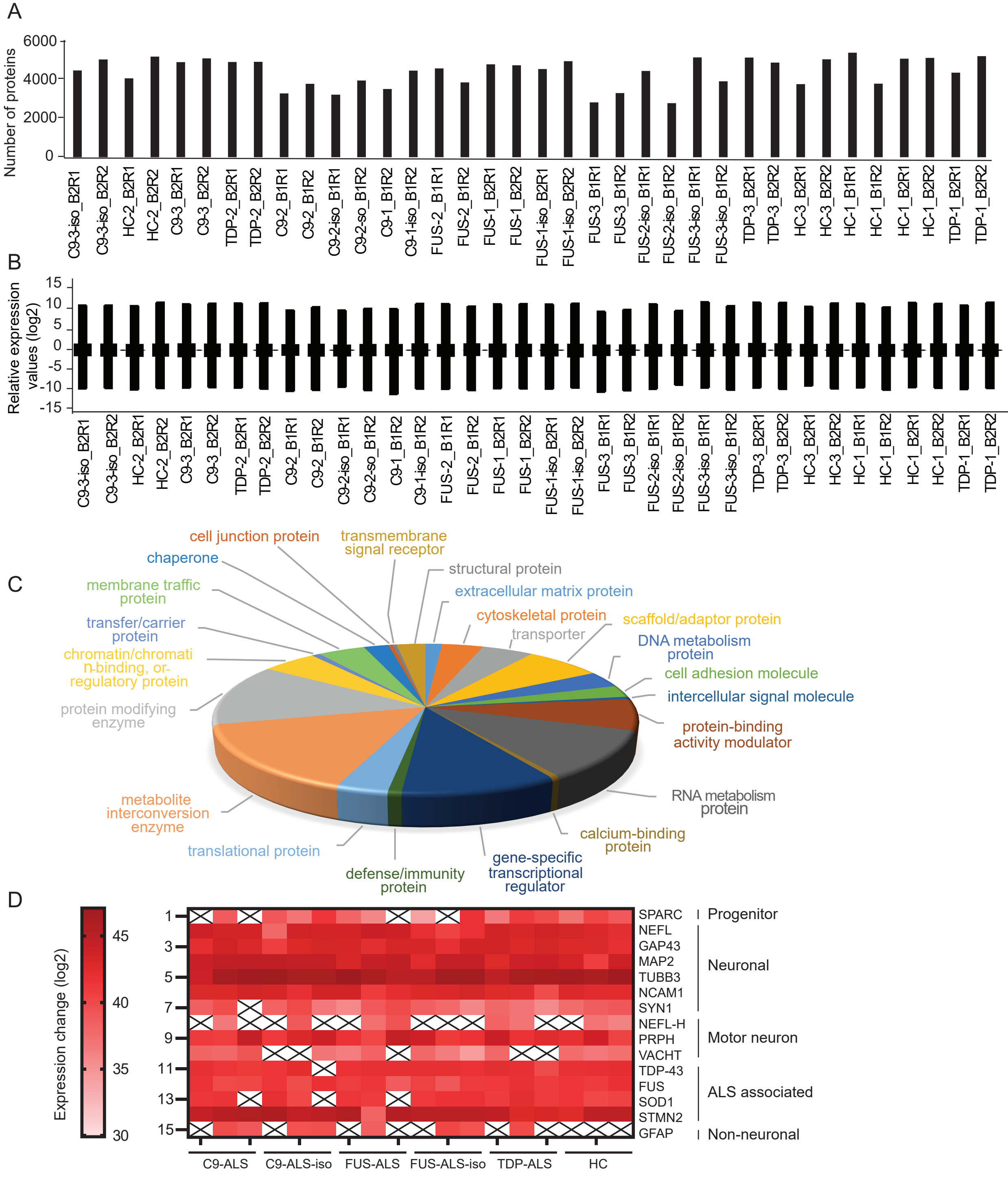

### Figure S3

#### Supplementary Figure 3

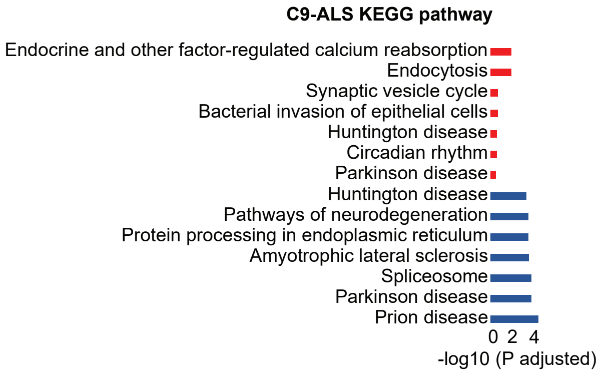


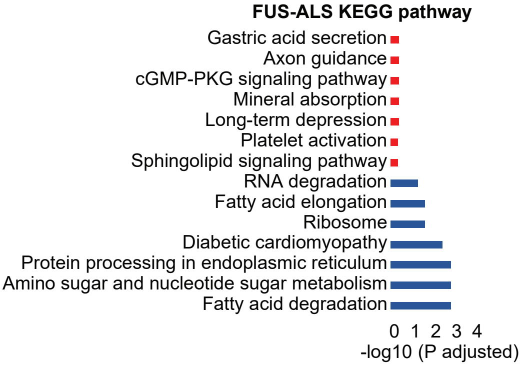


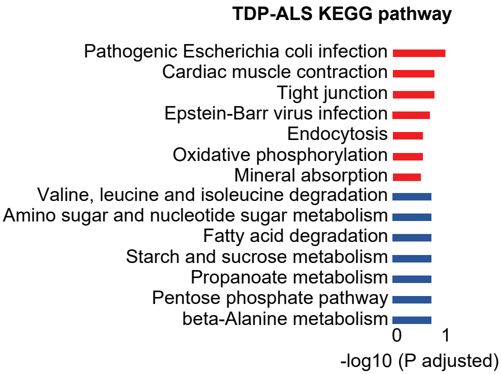

### Figure S4

#### Supplementary Figure 4

A

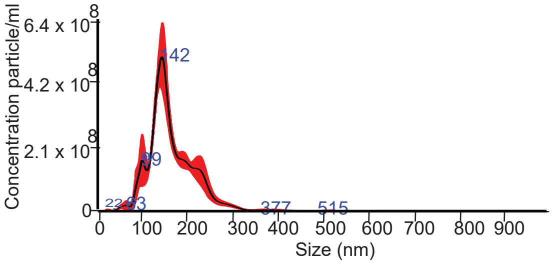


B

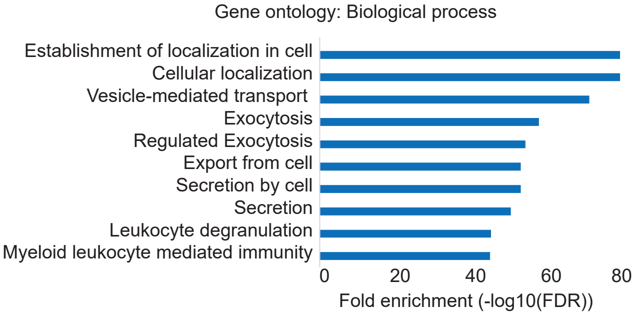


C

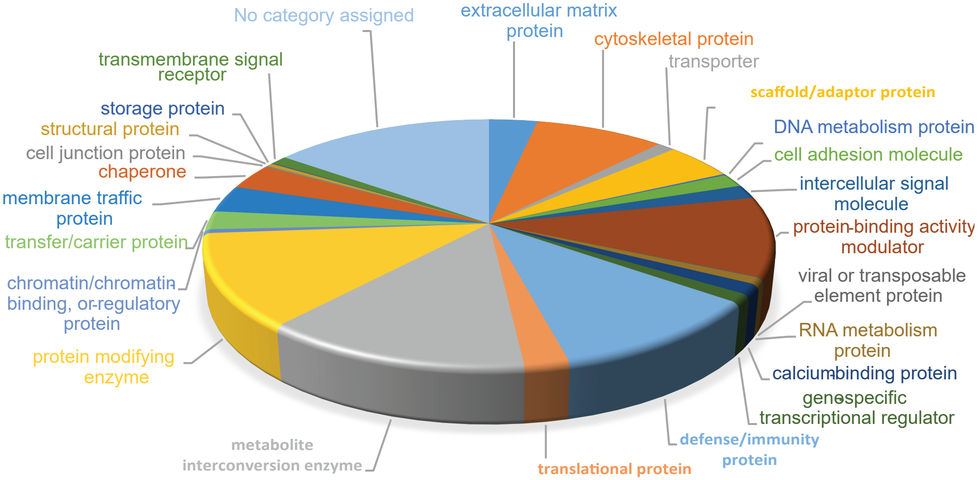


D

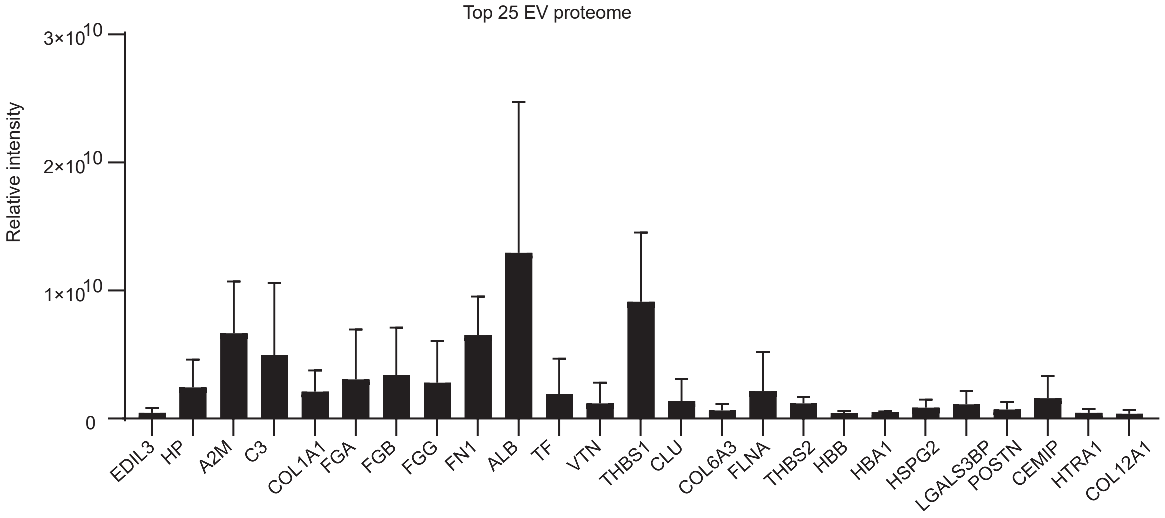
