## Supplementary material for "Proteomic signature and mesenchymal stromal/stem cell-derived extracellular vesicle treatment of iPSC motor neurons from different ALS patient groups": Table S1

**Supplementary Table 1.** Information of patient and control lines used for this study

| Cell line | Name in lab | Gender | Age of Biopsy | Virus used | Genotype | Where made | Reference | ALS pathology in iPSC-motor neurons |
| --- | --- | --- | --- | --- | --- | --- | --- | --- |
| C9ORF72 |  |  |  |  |  |  |  |  |
| C9-1 | CS52iALS-C9n6 | Male | 45 | Episomal Plasmid | C9orf72 Hexanucleotide repeat expansion (6-8kb) | Cedars Sinai | https://biomanufacturing.cedars-sinai.org/product/cs52ials-c9nxx/ ; Sareen et al.  (Science Translational Medicine, 2013) | RNA foci selectively in C9-ALS motor neurons; Increased DPRs and disrupted nucleocytoplasmic transport |
| C9-1-iso | CS52iALS-C9n6.ISOC3 | Male | 46 | Used CRISPR/Cas9 | C9 mutation corrected via CRISPR Cas9 | Cedars Sinai | https://biomanufacturing.cedars-sinai.org/product/cs52ials-c9n6-isoxx/ |  |
| C9-2 | CS29iALS-C9n1 | Male | 47 | Episomal Plasmid | C9orf72 Hexanucleotide repeat expansion (6-8kb expanded allele) | Cedars Sinai | https://biomanufacturing.cedars-sinai.org/product/cs29ials-c9nxx/ ;  Sareen et al. (Science Translational Medicine, 2013) | RNA foci selectively in C9-ALS motor neurons; Increased DPRs and disrupted nucleocytoplasmic transport |
| C9-2-iso | CS29iALS-C9n1.ISOT2RB4 | Male | 47 | Used CRISPR/Cas9 | C9 mutation corrected via CRISPR Cas9 | Cedars Sinai | https://biomanufacturing.cedars-sinai.org/product/cs29ials-c9n1-isoxx/ |  |
| C9-3 | C902-02 | Female | 51 | retrovirus | C9orf72 (clone 2 from patient C902) | Talbot Lab Oxford | <https://doi.org/10.1002/stem.2388> | Ca^2+^ dysregulation, ER stress, loss of proteostasis. Impaired mitochondrial Ca ^2+^ uptake and glutamate excitotoxicity |
| C9-3-iso | 59.1 (ISO_CTRL C902-02) | Female | 51 | Used CRISPR/Cas9 | C9orf72 most repeats deleted but locus intact (2 repeats left). | Talbot Lab Oxford | <https://doi.org/10.1002/stem.2388> |  |
| Healthy controls |  |  |  |  |  |  |  |  |
| HC-1 | PZ1-5 | Male | 62 | Sendai virus | Healthy ctrl | UMC Utrecht | https://doi.org/10.1186/s40478-024-01852-6 |  |
| HC-2 | 929c4 | Female | 60 | Sendai virus | Healthy ctrl | UMC Utrecht | made at UMC Utrecht Brain Center MIND facility  (manuscript in submission) |  |
| HC-3 | OH3.1 | Male | 49 | Lentivirus | Healthy ctrl | UMC Utrecht | 10.1038/s41467-018-06684-2 (iPSC 5) |  |
| TDP-43 |  |  |  |  |  |  |  |  |
| TDP-1 | TDP0102-M337V | Male | 61 | Sendai | TDP43 I383T mutation | University of Oxford | [10.1016/j.stemcr.2020.03.023](https://doi.org/10.1016/j.stemcr.2020.03.023) | High glutamate-induced Ca ^2+^ release, impaired mitochondrial Ca ^2+^ uptake and glutamate excitotoxicity |
| TDP-2 | CiRA 00024 | Female | 60 | retrovirus | TDP43 M337V mutation | Riken Japan | 10.2169/internalmedicine.49.2915; [DOI: 10.1126/scitranslmed.3004052](https://doi.org/10.1126/scitranslmed.3004052) | Cytoplasmic aggregates, shorter neurites |
| TDP-3 | M337V.2 | Male | 59 | Episomal reprogramming | TDP43 M337V mutation | University of Edinburgh | <https://doi.org/10.1073/pnas.1202922109> | Cytoplasmic accumulation of TDP-43, increased cell death |
| FUS |  |  |  |  |  |  |  |  |
| FUS-1 | FUS2 | Male | 26 | Lentivirus | R495QfsX527 Frameshift mutation | Hannover Germany | <https://doi.org/10.1002/stem.2354> | Hypoexcitability in iPSC-motor neurons, ion channel imbalances with lower sodium to potassium (Na^+^/K^+^) ratios |
| FUS-1-iso | FUS2 isogenic control | Male | 26 | CRISPR/Cas9 | WT FUS | Hannover Germany | <https://doi.org/10.1002/stem.2354> |  |
| FUS-2 | FUS 2/2 | Female | 71 | Sendai virus | R521H FUS Heterozygous point mutation | Leuven | [10.1038/s41467-017-00911-y](https://dx.doi.org/10.1038%2Fs41467-017-00911-y) | Cytoplasmic accumulation of FUS, hypoexcitability, mitochondrial and ER vesicle axonal transport defects. |
| FUS-2-iso | FUS CO2/2 | Female | 71 | Made from FUS2/2, company Cell System | FUS R521H | Cell System | [10.1038/s41467-018-06111-6](https://dx.doi.org/10.1038%2Fs41467-018-06111-6) |  |
| FUS-3 | FUS 3/1 | Male | 17 | Sendai virus | P525L FUS Heterozygous point mutation | Leuven | [10.1038/s41467-017-00911-y](https://dx.doi.org/10.1038%2Fs41467-017-00911-y) | Cytoplasmic accumulation of FUS, hypoexcitability, mitochondrial and ER vesicle axonal transport defects. |
| FUS-3-iso | FUS CO3/3 | Male | 17 | Made from FUS3/3, company Cell System | FUS P525 | Cell System | [10.1038/s41467-018-06111-6](https://dx.doi.org/10.1038%2Fs41467-018-06111-6) |  |
